# RORγt^+^CD4^+^ T cells promote IL-23R-mediated neuronal cell apoptosis in the central nervous system

**DOI:** 10.1101/2023.04.17.537133

**Authors:** Sandip Ashok Sonar, Thoihen Heikrujam Metei, Shrirang Inamdar, Nibedita Lenka, Girdhari Lal

## Abstract

Transcription factors T-bet and RORγt play a crucial role in neuronal autoimmunity, and mice deficient in these two factors do not develop experimental autoimmune encephalomyelitis (EAE). The independent role of T-bet and RORγt in the pathogenesis of EAE and how they help induce apoptosis of neurons in the central nervous system (CNS) during neuronal autoimmunity is unclear. In the present study, we showed that myelin oligodendrocyte glycoprotein (MOG_35-55_) peptide-specific Th1 cells deficient in RORγt could cross BBB but fail to induce apoptosis of neurons and EAE. Pathogenic Th17 cell-derived cytokines GM-CSF, TNF-α, IL-17A, and IL-21 significantly increase the surface expression of IL-23R on neuronal cells. Furthermore, we showed that, in EAE, neurons in the brain and spinal cord express IL-23R. IL-23-IL-23R signaling in neuronal cells caused phosphorylation of STAT3 (Ser727 and Tyr705) and induced cleaved caspase 3 and cleaved poly (ADP-ribose) polymerase-1 (PARP-1) molecules in an IL-23R-dependent manner and caused apoptosis. Thus, we provided a mechanism where we showed that T-bet is required to recruit pathogenic Th17 cells and RORγt expression to drive the apoptosis of IL-23R^+^ neurons in the CNS and cause EAE. Understanding detailed molecular mechanisms will help to design better strategies to control neuroinflammation and autoimmunity.

**GRAPHICAL ABSTRACT:** 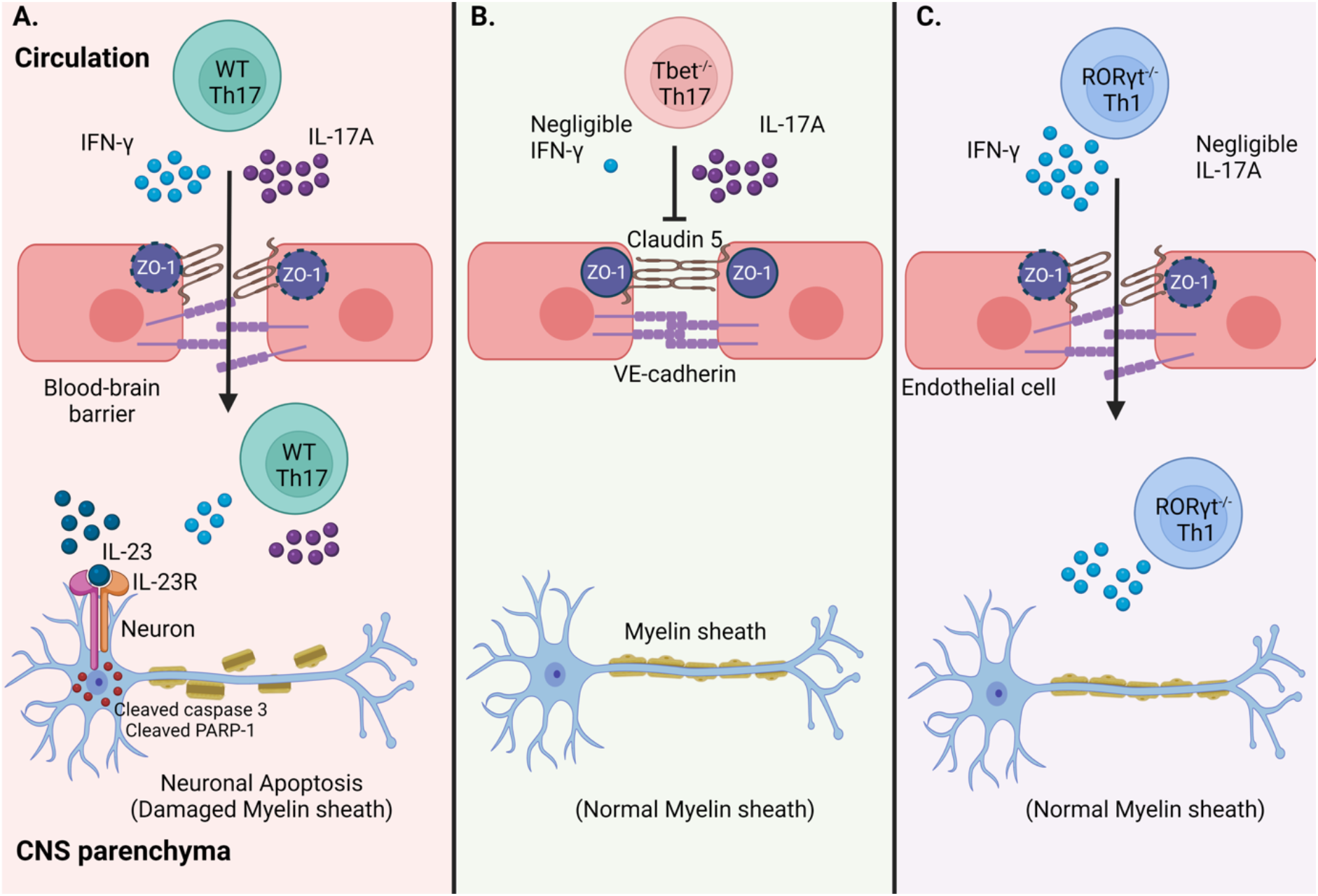

**One Sentence Summary:** IL-23-IL-23R signaling promotes apoptosis of CNS neurons.

## Introduction

The autoreactive Th1 and Th17 cells play a critical role in the pathogenesis of neuroinflammation and autoimmunity. Mice lacking transcription factors T-bet or RORγt do not develop experimental autoimmune encephalomyelitis (EAE) (*1*). T-bet expression and cytokine IFN-γ play a critical role in the pathogenicity of Th17 cells (*1–3*). It has been shown that T-bet^+^RORγt^+^ Th17 cells are more pathogenic than conventional Th17 cells (*4, 5*). The development of EAE requires a complex cellular and molecular interaction in peripheral lymphoid tissues and the central nervous system (*6*). The first step for CD4 T cell-mediated neuronal autoimmunity requires crossing CD4^+^ T cells to the blood-brain barrier (BBB) and then interacting these cells with CNS resident cells and neurons. Adoptive transfer of only neuronal antigen-specific Th1 or Th17 cells and peripheral inflammation in mice leads to the development of EAE (*7*). The ratio of Th17:Th1 cells dictates the location of CNS inflammation in the brain (*8*). The absence of peripheral inflammatory signals ablates myelin-specific CD4^+^ T cells to cause EAE (*9*). However, the independent role of specific transcription factors, T-bet and RORγt, in the transmigration of antigen-reactive CD4^+^ T cells into the CNS or the mechanism of inducing apoptosis of neurons in the CNS tissue is not clearly understood.

The RORγt is a well-known master regulator of Th17 cell differentiation and regulates the expression of several inflammatory molecules such as IL-17A, IL-17F, IL-21, IL-21R, IL-22, IL-23R, and GM-CSF in Th17 cells (*10, 11*). The signaling from IL-17A/F, IL-21, and IL-22 is redundant in EAE (*12–14*). However, IL-23R and GM-CSF signaling are pathologically important, and mice without IL-23R, IL-23, or GM-CSF are completely protected from the EAE (*15, 16*). The function of IL-23-IL-23R signaling is known to fine-tune the Th17 cells and confer a stable and highly pathogenic role (*15, 17*). The IL-23p19^-/-^ or IL-23R^-/-^ mice prevent clinical EAE development and are mainly attributed to the defect in the Th17 pathogenic role (*17, 18*). However, whether IL-23 signaling works at the level of CNS parenchyma is not addressed adequately. Moreover, does CNS-resident cells and neurons express IL-23R and respond to IL-23 stimulus during EAE is not well characterized.

In the present work, we show that T-bet controls the transmigration potential, whereas RORγt expression in CD4^+^ T cells drives the apoptosis of neurons leading to the development of EAE. Th17-specific cytokines GM-CSF, TNF-α, IL-17A, and IL-21 promote IL-23R expression in neuronal cells. Further, In EAE, neurons in the brain and spinal cord express IL-23R, and interaction with IL-23 causes apoptosis of neurons via activation of caspase 3 (cleavage of caspase 3) and cleavage of poly(ADP-ribose) polymerase-1 (PARP-1). The knockdown of IL-23R in neurons protected from the IL-23-mediated apoptosis. The present study identifies the spatiotemporal role of transcription factors T-bet and RORγt in the development and progression of EAE and the mechanism of neuronal death in EAE.

## Results

### RORγt^-/-^ Th1 cells migrate into CNS but fail to induce EAE

To dissect the contribution of T-bet and RORγt in the transmigration of pathogenic Th1 and Th17 cells and CNS pathology, we adoptively transferred *in vitro*-differentiated MOG_35-55_ (MOG)-specific wild-type Th17, T-bet^-/-^ Th17, or RORγt^-/-^ Th1 cells (all express CD45.2 marker) into recipient T-bet^+/+^ mice (CD45.1 congenic) **(Figure S1A)**, and the development of EAE was monitored. Our results showed that the adoptive transfer of only T-bet^+/+^ Th17 cells could cause demyelination and induce EAE but not T-bet^-/-^ Th17 cells or RORγt^-/-^ Th1 cells **(Figures S1B and 1A)**. Furthermore, the mice that received T-bet^-/-^ Th17 cells showed reduced or no infiltration of CD45^+^ leukocytes or CD4^+^ T cells in the brain and spinal cord compared to recipients of T-bet^+/+^ Th17 cells **(Figure 1B)**. Although the adoptive transfer of RORγt^-/-^ Th1 or T-bet^-/-^ Th17 did not cause EAE, RORγt^-/-^ Th1 recipients showed increased infiltration of total CD45^+^ leukocytes as well as CD4^+^ T cells as compared to the T-bet^-/-^ Th17 group **(Figure 1B)**. The migration of CD45^+^ leukocytes and CD4^+^ T cells in the recipients of RORγt^-/-^ Th1 cells was comparable to the recipients of T-bet^+/+^ Th17 cells in the brain but significantly reduced numbers of cells found in the spinal cord **(Figure 1B)**.

**Figure 1.**
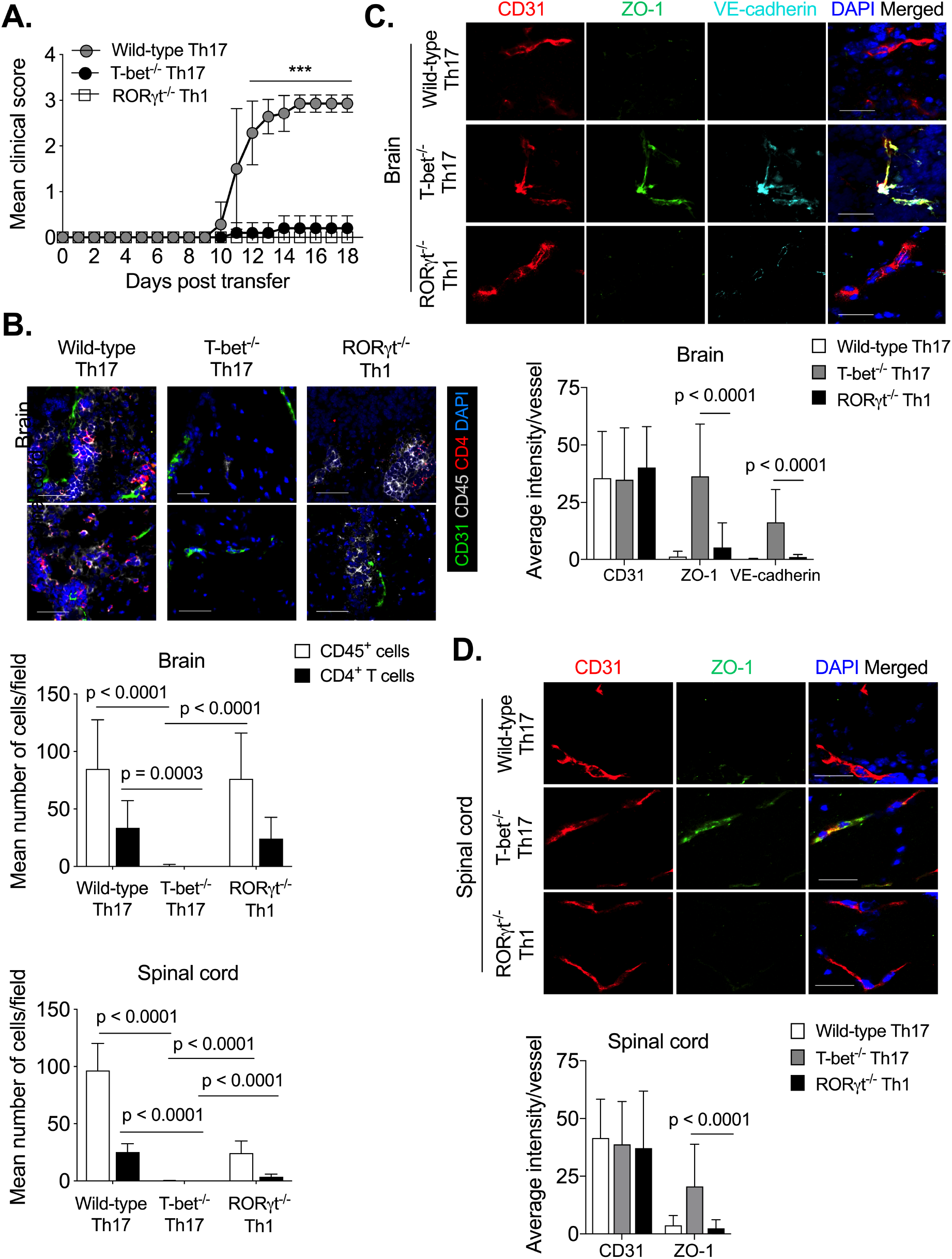
T-bet^-/-^ Th17 failed, but RORγt^-/-^ Th1 cells damage the BBB, and both cells do not induce disease. MOG-specific Th17 cells from wild-type (CD45.2) and T-bet^-/-^ (CD45.2) mice or Th1 cells from RORγt^-/-^ (CD45.2) mice were generated by *in vitro* culture of CD4^+^ T cells. Th1 or Th17 cells (20 x 10^6^ cells/mouse) were adoptively transferred into wild-type (CD45.1 congenic) mice and given PTx on days 0 and 2. **(A)** Mean clinical score of EAE plotted. n = 6-7 mice/group. **(B)** On day 18, brain and spinal cord tissues were stained with CD4^+^ (red), leukocyte marker CD45 (grey), CD31^+^ endothelial cells (green), and DAPI (dark blue). The representative images are shown (upper). CD4^+^ and CD45^+^ cells/field were quantitated. A total of 12-42 images from the brain and spinal cord sections were quantified and plotted (lower). (n=3-5 mice/group). **(C, D)** On day 18, the brain and spinal cord were stained with CD31^+^ endothelial cells (red), ZO-1 (green), VE-cadherin (light blue), and nuclear stain DAPI (dark blue), and a representative image is shown (upper). Quantitation of the average intensity of CD31, ZO-1, and VE-cadherin/vessel is shown (lower). 27 to 39 vessels in the brain and 26 to 64 vessels in the spinal cord were analyzed. Data are representative of two independent experiments with 3-5 mice brain and spinal cord tissues from each group **(B- D)**; *** p < 0.0001; One-way ANOVA followed by Tukey’s test **(A-D)**; Original magnification 400x **(B)**, 630x **(C, D)**; Error bar represents the standard deviation (S.D.) **(A-D)**; Scale bar 30μm **(C, D)** and 50μm **(B)**.

The BBB endothelium is a significant physiological barrier separating peripheral circulation from the CNS parenchyma (*19*). The BBB endothelial junctions are very dynamic and form zipper-like structures sealing the BBB. This restrictive barrier is critical for maintaining CNS homeostasis and selectively allows nutrient transport, and restricts inflammatory mediators and immune cells’ entry into the CNS parenchyma (*19*). Pathogenic CD4^+^ T cells must breach this barrier to enter the CNS parenchyma and cause EAE (*6, 20*). In our experiments, the immunohistological analysis of the brain and spinal cord in EAE mice showed that T-bet^+/+^ Th17 and RORγt^-/-^ Th1 transferred groups have damaged BBB endothelium junctions, whereas T-bet^-/-^ Th17 group did not show similar changes **(Figures 1C, 1D, and S1C)**. Interestingly, although the RORγt^-/-^ Th1 transferred group had disrupted BBB in the spinal cord, fewer leukocytes and CD4^+^ T cells could enter the spinal cord **(Figure 1B, 1D, and S1C)**. These results suggest that encephalitogenic Th17 cells require T-bet to alter the adherence and tight junction molecules at the BBB, and RORγt^-/-^ Th1 cells that possess functional T-bet expression and are competent to produce the IFN-γ able to breach the BBB.

### RORγt^-/-^ Th1 cells failed to induce iNOS in the host astrocytes in the CNS potently

Recent evidence indicates that astrocytes play protective and detrimental roles in early MS lesion formation and disease progression (Ponath et al., 2018; Ponath et al., 2017). During EAE, reactive astrocytes secrete nitric oxide, which mediates axonal damage during neuroinflammation (*21, 22*). In the CNS, various extracellular signals induce the expression of inducible nitric oxide synthase (iNOS) in astrocytes, and IFN-γ is a primary inducer of iNOS production in neutrophils, microglial cells, and astrocytes and causes apoptosis in the infiltrating immune cells (*23*). We examined whether the inability of RORγt^-/-^ Th1 cells to induce EAE is due to apoptosis of these cells in the inflamed CNS or lack of Th17-associated factors. Adoptive transfer of wild-type Th17 cells showed increased iNOS expression in the astrocytes (GFAP^+^ cells) in the spinal cord **(Figure 2A)** and increased apoptosis (cleaved caspase 3^+^ cells) in the oligodendrocytes (CNPase^+^ cells; myelin-producing cells in the CNS) and to some extent in the infiltrating leukocytes (CD45^+^ cells) in the brain and spinal cord **(Figure 2B)**. Adoptive transfer of RORγt^-/-^ Th1 cells poorly induced the astrocytic iNOS expression **(Figure 2A)** and showed a significantly reduced number of oligodendrocytes with apoptotic cleaved caspase 3 stainings **(Figure 2B)**. Interestingly, these fewer iNOS^+^ astrocytes seemed to cause the apoptosis of an increased number of leukocytes **(Figures 2A and 2B)**. T-bet^-/-^ Th17 cells did not show infiltration of leukocytes, iNOS production, or apoptosis in the brain and spinal cords **(Figures 2A and 2B)**. To test the role of iNOS induced-apoptosis of infiltrating immune cells in the recipients of RORγt^-/-^ Th1 cells, we generated MOG-specific wild-type Th1, and RORγt^-/-^ Th1 cells and adoptively transferred them in iNOS^-/-^ mice. Our results showed that wild-type Th1 cells induced EAE symptoms in iNOS^-/-^ mice **(Figure 2C)** and showed increased infiltration of CD4^+^ T cells and CD45^+^ leukocytes in the brain and spinal cord (**Figure 2D)**. However, RORγt^-/-^ Th1 cells failed to induce EAE in iNOS^-/-^ recipients **(Figure 2C)** and had reduced infiltration of leucocytes and CD4^+^ T cells in the CNS compared to the wild-type Th1 recipient group (**Figure 2D)**. These results suggest that although iNOS induces apoptosis in the infiltrating inflammatory cells in the RORγt^-/-^ Th1 transferred group, iNOS is dispensable for EAE induction.

**Figure 2.**
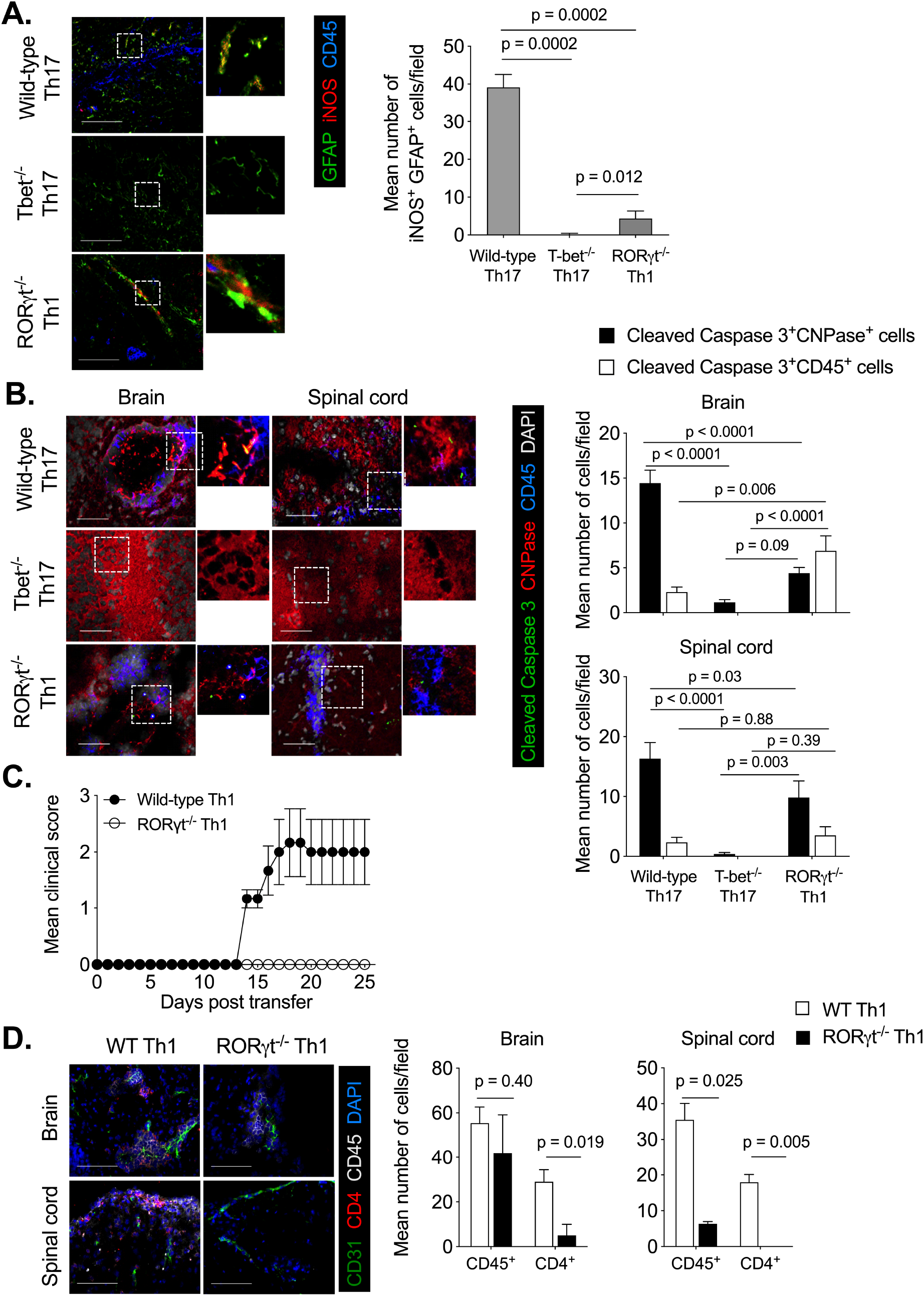
Host-derived iNOS is dispensable to induce apoptosis in leukocytes in the CNS. At the peak of the disease (day 18), spinal cord tissues of mice from figure 1A were stained for (A) astrocyte marker GFAP (green), iNOS (red), and leukocyte marker CD45 (blue), and representative images are shown (left). The dotted white square marked in the image is magnified and shown on the right. The number of GFAP^+^ cells positive for iNOS staining was counted and plotted as the mean number of cells/field (right). **(B)** Brain and spinal cord tissues were stained with apoptotic cell marker cleaved caspase 3 (green), oligodendrocyte marker CNPase (red), and leukocyte marker CD45 (dark blue), and nuclear stain DAPI (grey), and a representative image is shown (left). The dotted white square marked in the image is magnified and shown on the right. The number of cells positive for the stain was counted and plotted as the mean number of cells/field (right). Data represent 3-5 mice brain and spinal cord tissues from each group. **(C)** MOG-specific Th1 cells from wild-type and RORγt^-/-^ mice were generated by *in vitro* culture of CD4^+^ T cells. Th1 cells (10 x 10^6^ cells/mouse) were adoptively transferred into iNOS^-/-^ mice and given PTx on days 0 and 2. Mean clinical scores of recipient iNOS^-/-^ mice were plotted. **(D)** Mice from Figure 3C were sacrificed on day 25, and the brain and spinal cord were analyzed by immunofluorescence staining. Representative images stained with CD4 (red), CD45 (grey), CD31 (green), and nuclear stain DAPI (blue) were shown (left). The mean number of CD45^+^ and CD4^+^ cellular infiltrations from 15-20 images of the brain and spinal cord is quantitated and plotted (right). Mann-Whitney test **(A)**, two-way ANOVA followed by Tukey’s test **(B),** and Student t-test **(D)**. Original magnification 400x **(A, D)** 630x **(B)**, scale bar 50 μm **(B)** and 100 μm **(A, D)**.

### RORγt^-/-^ Th1 cells fail to induce IL-23 receptors on CNS neurons

Since our results showed that iNOS-induced apoptosis of infiltrating immune cells is dispensable for EAE induction in the RORγt^-/-^ Th1 transferred group, we hypothesized that the lack of clinical EAE might be due to the lack of other RORγt-dependent pathogenic factors in the inflamed CNS microenvironment. RORγt-dependent transcriptional program promotes the expression of IL-17A, IL-21, IL-22, GM-CSF, and IL-23R molecules, and these molecules regulate the inflammatory response in the CNS (*11, 18, 24*). For example, IL-17RA signaling promotes CNS inflammation through glial cell populations, specifically NG2^+^ glial cells (*25, 26*). However, several studies with gene deletion mouse strains have revealed the redundant roles of IL-17A, IL-17F, IL-21, and IL-22 in promoting CNS inflammation (*12, 14, 27*). Nonetheless, GM-CSF and IL-23 (p19 subunit) have a non-redundant pathogenic role in neuroinflammation and autoimmunity (*11, 28, 29*). Most studies have linked the pathogenic role of IL-23 with the generation and maintenance of highly pathogenic Th17 cells. Recently, the development of a transgenic mouse model over-expressing astrocyte-restricted IL-23 showed a variety of neuroinflammatory phenotypes, including exacerbated EAE symptoms (*30, 31*). Neutralization or deficiency of GM-CSF, IL-23 (p19 subunit), or IL-23R prevents the demyelination of CNS tissues and inhibits active EAE (*11, 29, 32, 33*). However, very little is known about the role of IL-23R signaling in CNS resident cells and neurons in EAE.

To understand the role of IL-23-induced signaling in regulating CNS pathology in MOG-specific RORγt^-/-^ Th1 recipients, we tested the expression of IL-23R in the brain and spinal cord. The immunohistochemical analysis of the brain and spinal cord of EAE mice showed the expression of IL-23R-expressing β-III tubulin^+^ neurons **(Figure 3A)**. Furthermore, we did not detect IL-23R expression on CNPase^+^ oligodendrocytes **(Figure 3B)** or F4/80^+^ microglia/macrophages **(Figure 3C)** in mice having EAE. Further, mice that received MOG-specific wild-type Th17 cells had a significantly higher number of IL-23R-expressing NeuN^+^ neurons in the brain than the RORγt^-/-^ Th1 cells group **(Figure 3D)**. The T-bet^-/-^ Th17 cell recipients showed a complete absence of IL-23R on neurons **(Figure 3D)**. Interestingly, we found that RORγt^-/-^ Th1 cell recipients completely lacked the presence of IL-23R^+^CD11b^+^ cells in the brain but showed significantly lower IL-23R^+^CD11b^+^ cells as compared to the wild-type Th17 group in the spinal cord **(Figure 3E)**. Furthermore, we did not find either CD11b^+^ cells or IL-23R^+^CD11b^+^ cells in T-bet^-/-^ Th17 transferred group **(Figure 3E)**. Together, these results showed that during EAE, neurons express IL-23R, and wild-type (T-bet^+/+^) MOG-specific Th17 cell-induced inflammation promotes IL-23R expression on NeuN^+^ neurons. In contrast, RORγt^-/-^ Th1 cell-induced inflammation failed to induce IL-23R expression on the neuron.

**Figure 3.**
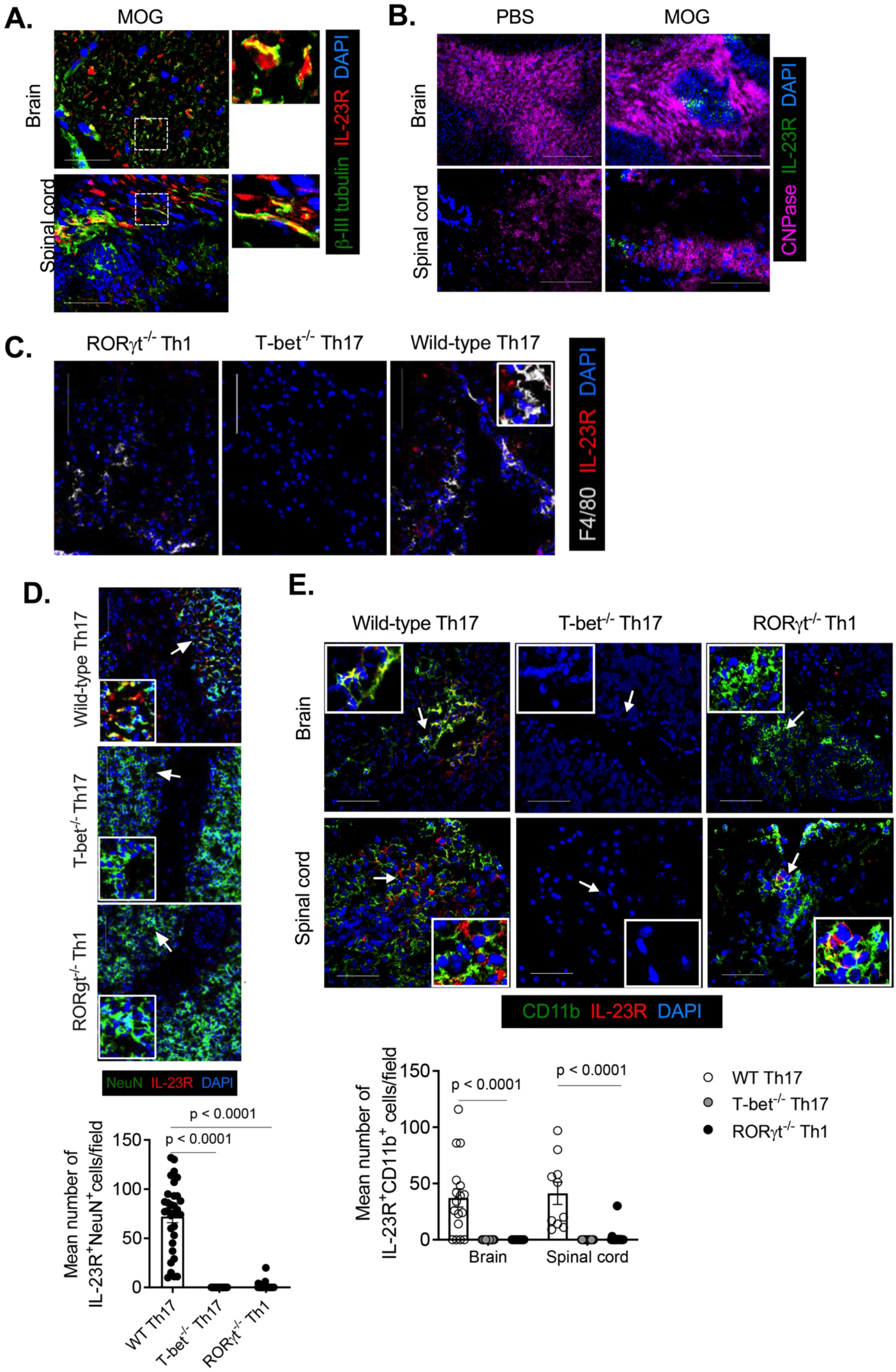
IL-23R expression in CNS during EAE. Active EAE was induced in C57BL/6 mice by *s.c.* injection of MOG and *i.v.* injection of PTx at days 0 and 2. Mice were sacrificed on day 21 (peak of disease), and the brain and spinal cord tissue sections were stained for **(A)** β-III tubulin (green), IL-23R (red), and nuclear stain DAPI (dark blue). **(B)** CNPase (magenta), IL-23R (green), and nuclear stain DAPI (dark blue). Representative images are shown. The regions marked with the dotted square are shown on the right side. **(C)** The brain tissues of mice from Figure 1A were stained for F4/80 (grey), IL-23R (red), and nuclear stain DAPI (dark blue). **(D)** Brain stained for NeuN^+^ (green), IL-23R^+^ (red), and nuclear stain DAPI (dark blue), and the representative images are shown (left). The regions marked with white arrows are shown in the inset of each image. The mean number of IL-23R^+^NeuN^+^ cells was counted from different images and plotted (right). **(E)** CD11b (green), IL-23R (red), and nuclear stain DAPI (dark blue). The representative images are shown. The regions marked with white arrows are shown in the inset. Data are representative of two independent experiments. n=3-4 mice/group. Mann-Whitney test **(D).** Two-way ANOVA followed by Sidak’s multiple comparison tests **(E)**. Original magnification 100x **(B, D,** Brain), 400x **(A,** Spinal cord**; C, E,** Brain). Scale bar 50 μm **(E)**, 100 μm **(A, B,** Spinal cord), 300 μm **(D,** Brain).

### IL-23R^+^ neurons undergo apoptosis during EAE

To further understand the *in vivo* relevance of IL-23R signaling on neurons, we injected recombinant purified mouse IL-23 (mIL-23) via intracerebroventricular (*i.c.v.*) route in the MOG-immunized RORγt^+/-^ and RORγt^-/-^ mice and assessed the IL-23R expression on neurons, and analyzed its role in the apoptosis of neurons. Our results showed that a single *i.c.v.* injection of IL-23 induced the expression of IL-23R on β-III tubulin^+^ neurons in the spinal cord of RORγt^+/-^ mice, but it was significantly lowered in RORγt^-/-^ mice **(Figure 4A)**. Interestingly, IL-23R-expressing neurons (MAP2^+^ cells) showed a sign of apoptosis as assessed by the cleaved caspase 3^+^ cells **(Figure 4B)**. We also observed an increased number of CD11b^+^ microglia in the dentate gyrus region and CD11b^+^ monocytes/macrophages infiltration in the ventricular area of RORγt^+/-^ mice compared to RORγt^-/-^ mice **(Figure 4C)**. To understand the absence of EAE symptoms in the recipients of RORγt^-/-^ Th1 cells, we tested the effect of IL- 23R expression on the apoptosis of neurons. Our immunohistochemical analysis showed the presence of cleaved caspase 3^+^IL-23R^+^β-III tubulin^+^ neurons suggesting the apoptosis of neurons in the spinal cord of the MOG-specific wild-type Th17 cells recipient group **(Figure 4D)**. However, the recipients of T-bet^-/-^ Th17 and RORγt^-/-^ Th1 cells showed a significantly reduced number of cleaved caspase 3^+^IL-23R^+^β-III tubulin^+^ neurons **(Figure 4E)**. Collectively, these results suggest that, unlike T-bet^+/+^ Th17 cells, inflammation induced by RORγt^-/-^ Th1 cells failed to induce IL-23R expression on neurons, and therefore, lack of IL-23R-dependent apoptosis in neurons may contribute to the absence of neuronal damage and development of EAE in the recipients of RORγt^-/-^ Th1 cells.

**Figure 4.**
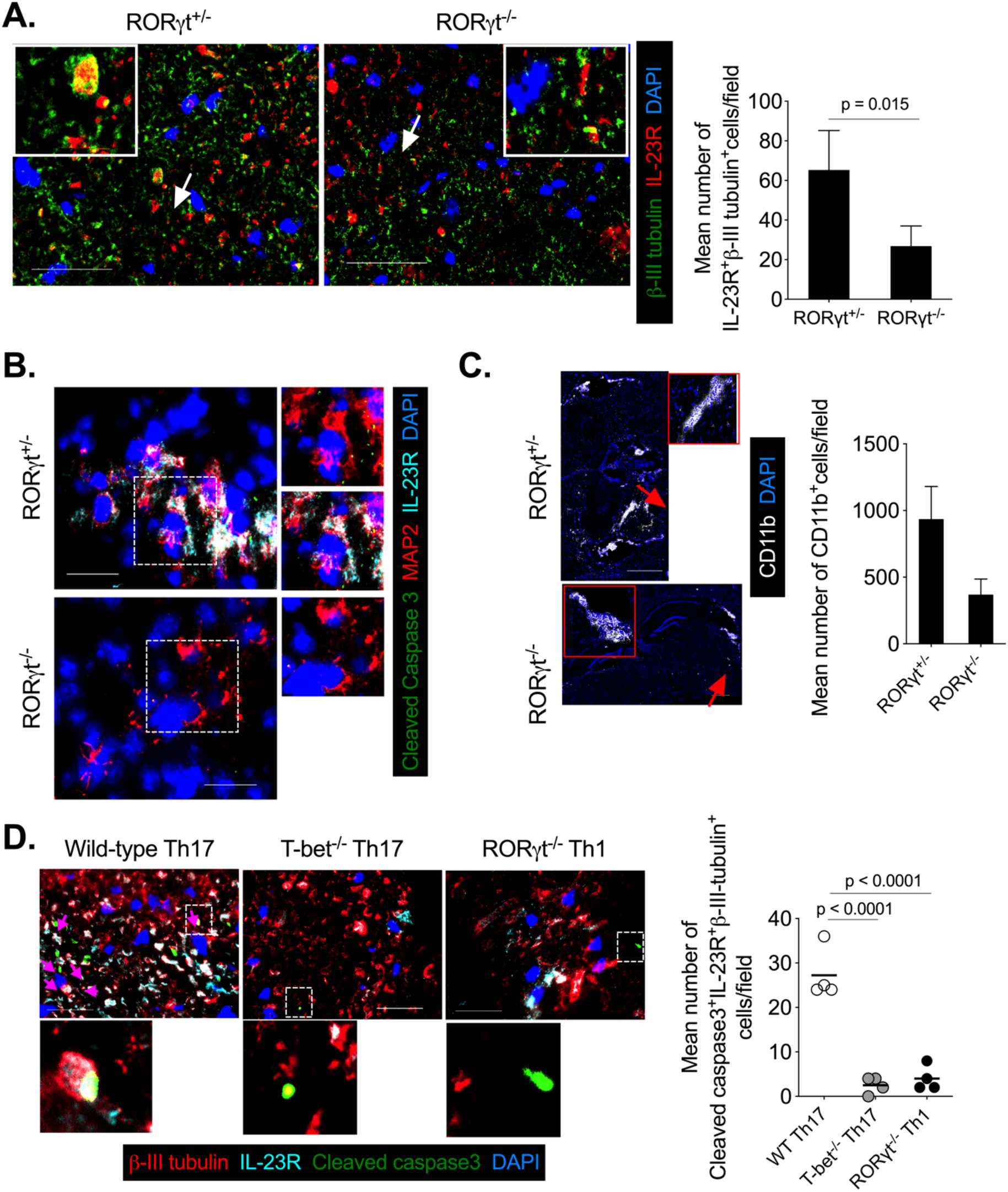
RORγt^-/-^ Th1 cells fail to induce IL-23R expression and cause apoptosis in neurons. At the peak of the EAE (day 18), RORγt^+/-^ or RORγt^-/-^ mice were stereotactically given intracerebroventricular (i.c.v.) injection of recombinant purified mouse IL-23 (200 pg/mouse). On day 3, the spinal cord was harvested, and sections were stained for **(A)** neuron- specific β-III tubulin (green), IL-23R (red), and nuclear stain DAPI (dark blue), and the representative images were shown (left). The regions marked with white arrows are shown in the inset of each image. The mean number of IL-23R^+^β-III tubulin^+^ cells was counted and plotted (right). **(B)** Brain sections were stained with MAP2 (red), IL-23R (light blue), cleaved caspase 3 (green), and nuclear stain DAPI (dark blue), and the representative images were shown. **(C)** Brain sections were stained for CD11b (grey) and nuclear stain DAPI (dark blue), and the representative images are shown (left). The regions marked with red arrows are shown in the inset of each image. The mean number of CD11b^+^ cells was counted and plotted (right). **(D)** The spinal cord tissues of mice from Figure 1A were stained for β-III tubulin (red), IL-23R (light blue), cleaved caspase 3 (green,) and nuclear stain DAPI (dark blue), and representative images are shown (left). The regions marked with dotted arrows are shown on the right side of each image. The mean number of cleaved caspase 3^+^IL-23R^+^β-III tubulin^+^ cells was counted from at least 10-15 different images from each mouse and plotted (right). The horizontal line denotes the mean, and each dot represents an individual mouse. Data represent two independent experiments with 3-4 mice brain and spinal cord tissue **(A-D)**. Mann-Whitney test **(B, D)**. Original magnification 100x **(C)**, 400x **(A)**, 1000x **(B, D)**. Scale bar 30 μm **(B, D)**, 100 μm **(A),** 600 μm **(C)**.

### IL-23-IL-23R signaling in neuronal cells causes STAT3-dependant apoptosis of neurons

To test that neuronal cells undergo IL-23-IL-23R-dependent apoptosis, we used a mouse neuroblastoma cell line (Neuro-2a) model for neuronal cells and analyzed IL-23R expression. Neuro-2a showed the mRNA and the surface expression of IL-23R **(Figure 5A)**. To understand the inflammatory cues capable of inducing IL-23R expression on neuronal cells during CNS inflammation, we tested the ability of individual cytokine treatments to change IL- 23R expression levels on Neuro-2a cells. The stimulation with inflammatory cytokines produced by pathogenic Th17 cells such as GM-CSF, TNF-α, IL-17A, IL-21, or cytokines required for the differentiation of Th17 cells such as IL-6 significantly increased the expression of IL-23R on neuronal cells **(Figure 5B and Figure S2)**, suggesting that the cellular sources of these cytokines are possible triggers of inducing IL-23R on neurons during EAE. Cytokines produced by Th1 cells, such as IFN-γ and IL-1β or cytokine required for Th1 cell differentiation IL-12 did not significantly alter the expression of IL-23R on neuronal cells **(Figure 5B** and **Figure S2)**. Further, to understand IL-23R signaling in neuronal cells, IL-23R was knockdown using IL-23R-shRNA **(Figure 5C)**. Stimulation of Neuro-2a cells with recombinant mouse IL-23 induced phosphorylation of signal transducer and activator of transcription (STAT3) specifically at Ser727 and Tyr705 residues in a time and IL-23 dose-dependent manner **(Figure 5D)**, suggesting that Neuro-2a cells express functional IL-23R. The importance of these site-specific activations of STAT3 by IL-23 in the neurons is largely unknown. The activation of caspase 3 occurs via the death ligand induced-cell extrinsic and mitochondrial intrinsic apoptosis pathway and plays a key role as an executor apoptotic cell death program (*34*). Interestingly, upon stimulation with recombinant IL-23, Neuro-2a cells showed an increased frequency of cleaved caspase 3, a marker for cells undergoing apoptosis in IL-23R^+^ neuronal cells **(Figures 5E** and **5F),** and were IL-23R dependent **(Figure 5F and 5G)**. Similar to cleaved caspase 3, the activation of poly(ADP-ribose) polymerase-1 (PARP-1) is also linked with the apoptosis process (*35*). PARP proteins catalyze the poly(ADP-ribosyl)ation of nuclear proteins and are also known to function in DNA damage repair processes. However, some apoptotic signals induce PARP-1 cleavage, rendering defects in DNA repair mechanisms. Important to note caspase 3 directly induces the cleavage of PARP-1 at Asp^214^ and Gly^215^, giving rise to 24- and 89-kDa fragments possessing DNA-binding (retained in the nucleus) and catalytic activity (retained in the cytosol), respectively (*36, 37*). PARP-1 cleavage inhibits DNA repair activity and favors the apoptosis program in the cell (*36*). We find that IL-23 increased cleaved PARP-1 in a time-dependent manner in neuronal cells (data not shown). Furthermore, to understand the importance of the expression of IL-23R on neurons, we *in vitro* differentiated mouse embryonic stem cells (ESCs) into the microtubule-associated protein 2 expressing (MAP2^+^) neuronal cells **(Figure S3A)**. These ESCs had a transgenic expression of tyrosine hydroxylase-GFP (Th. GFP) to track neural lineage cell differentiation. Flow cytometry analysis showed that Th.GFP^+^ neural cells showed surface expression of IL-23R **(Figure S3B)**. Stimulating these cultured neuronal cells with recombinant IL-23 led to increased apoptosis **(Figure S3C)**. Moreover, upon IL-23 stimulation, IL-23R-expressing GFP^+^ neurons showed an increased frequency of cleaved caspase 3 positive cells, an indicator of caspase 3 activation **(Figure S3D)**. These results suggest that neuronal cells express functional IL-23R on their surface, and IL-23 signaling induces the apoptotic program in the neurons.

**Figure 5.**
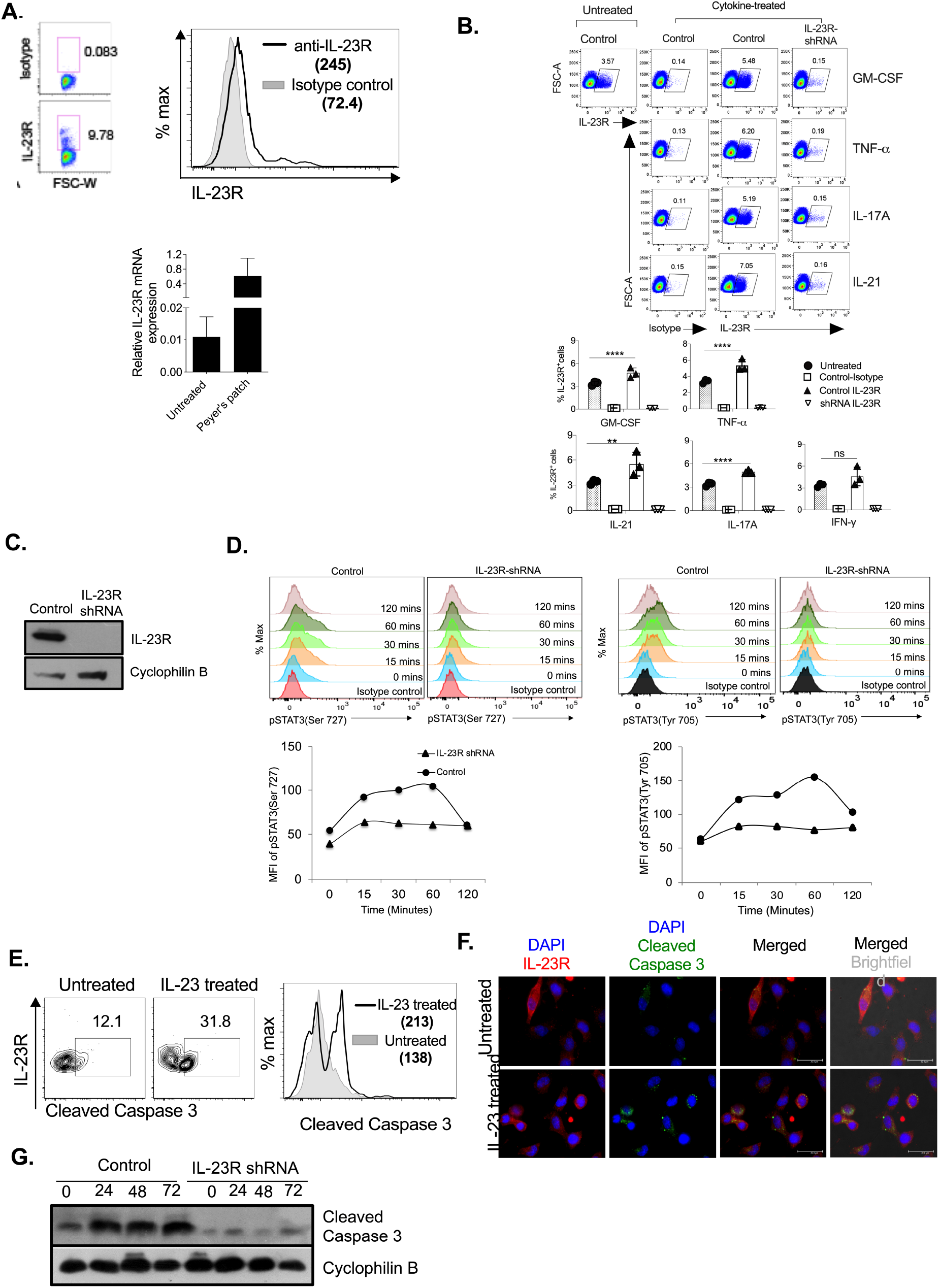
IL-23-IL-23R signaling induces apoptosis in neuronal cells. **(A)** Dot plots show surface IL-23R expression on Neuro-2a cells stained with anti-IL-23R antibody or isotype control antibody (Upper left panel). The number in the histogram (Upper right) represents MFI. Relative IL-23R mRNA expression in Neuro-2a cells and Peyer’s patch cells from wild-type C57BL/6 mice ( as a positive control) are shown (bottom panel). **(B)** Neuro-2a cells (control cells) or IL-23R knockdown Neuro 2a cells (IL-23R-shRNA) were treated with recombinant purified GM-CSF, TNF-α, IL-17A, IL-21, and IFN-γ (10 ng/ml) for 48 hours, and IL-23R expression were monitored using flow cytometry. A representative dot plot is shown (top). Each symbol represents data from an individual test. Statistical significance between control neuro2a cells treated and untreated samples are determined by One-way ANOVA. ** p <0.01; **** p < 0.0001; ns, not significant. **(C).** IL-23R was knockdown using shRNA. Expression of IL-23R in IL-23R knockdown cells is shown. **(D)** IL-23R knockdown or control Neuro-2a cells were treated with IL-23 (20 ng/ml) for indicated time points and stained for phosphoproteins- pSTAT3(Ser727) and pSTAT3(Tyr705). Histograms show the pSTAT3 (Ser727) and pSTAT3 (Tyr705) molecules in a time-dependent manner (top). MFI of pSTAT3 (Ser727) and pSTAT3(Tyr705) was calculated and plotted (bottom). **(E)** Dot plots show the expression of intracellular cleaved caspase 3 in IL-23-treated Neuro-2a cells for 48 hours. Data shown are gated on live, singlet, IL-23R^+^ cells (left). The histogram shows the expression of intracellular cleaved caspase 3 in cells gated on live, singlet, IL-23R^+^ neuro-2a cells. The numbers in the histogram indicate MFI. **(F)** Neuro-2a cells were treated with purified recombinant IL-23 (20 ng/ml) for 0, 24, 48, and 72 hours, and expression of cleaved caspase 3 was analyzed by Western blot. Cyclophilin B is used as the loading control. **(G)** The representative images of Neuro-2a cells stained with IL-23R (red), cleaved caspase 3 (green), and nuclear stain DAPI (blue) are shown. The data shown represent two **(A-C, F, G)** and three **(D, E)** independent experiments. Original magnification 1000x; scale bar, 30 μm **(G)**.

## Discussion

Activation of myelin-reactive T lymphocytes, their transmigration across the BBB into the CNS, and inflammatory damage in the CNS parenchyma orchestrate the pathology of MS in susceptible humans and EAE in rodents. Recent reports have suggested that the absence of T-bet or RORγt confers protection from EAE. This defect in the induction of EAE was explained due to the defect in the generation of autoreactive Th1 and Th17 cells in the periphery (*1, 3, 10*). Interestingly, in our study, adoptive transfer of MOG-specific Th17 lacking T-bet or Th1 cells lacking RORγt failed to induce EAE in wild-type recipients, indicating that these cells lacked encephalitogenic potential. The inability of these cells to induce EAE might be due to the following reasons- i) defect in the generation of effector cells in the periphery, ii) defective TEM across the BBB, and iii) defect in the induction of inflammation in the CNS parenchymal region. In this study, we also showed that T-bet^-/-^ Th17 cells had an increased frequency of IL-17A-expressing cells but lacked IFN-γ^+^ and had a low frequency of IL- 17A^+^IFN-γ^+^ to wild-type Th17 cells. Interestingly, unlike wild-type MOG-specific Th17 cells, T-bet^-/-^ Th17 failed to induce the alterations in the junctional molecules of the BBB, and we speculate that this might have resulted in the lack of TEM in the CNS. Recently, using *in vitro* BBB models, we showed that T-bet downstream target, IFN-γ acts on brain endothelial cells, affect the distribution of claudin-5, VE-cadherin, and ZO-1 molecules, and promote the apical to the basal side TEM of CD4^+^ T cells in an ICAM-1 and STAT-1-dependent manner (*38*). It has been shown that Th1-like Th17 (IFN-γ^+^IL-17A^+^ or T-bet^+^RORγt^+^) and ex-Th17 Th1 (IFN-γ^+^T-bet^+^ derived from T-bet^+^RORγt^+^ population) cells are abundantly present in the inflamed CNS (*39, 40*). These cells are highly encephalitogenic and show T-bet expression (*2, 41*). Although our MOG-specific T-bet^-/-^ Th17 cells were generated in the presence of IL-23, they lacked T-bet expression, and, therefore, the possibility of generation of pathogenic Th1-like

Th17 and ex-Th17 Th1 cells would be negligible. Earlier studies on T-bet’s role in encephalitogenic Th17 function have yielded mixed results. While some studies showed its essential role in the process, others documented that it is dispensable in pathogenic Th17 generation and EAE induction (*1, 3*). The discrepancies in these reports may be due to the differences in protocols employed to generate Th17 cells, methodologies used to knock down or knock out the T-bet gene, and different transgenic and knockout mouse lines used in the study. Moreover, though highly purified Th17 cells lacking T-bet showed elevated IL-17A and negligible IFN-γ expression, these cells recovered from the inflamed CNS exhibited considerable proportions of IL-17A^+^IFN-γ^+^ cells (*42–44*). Additionally, MOG-specific Th17 cells are highly heterogeneous, with varying cytokine secretion capability and different transcription factor expression. Our results showed that T-bet deficiency hampers the migration of Th17 cells in the CNS, affecting their encephalitogenic function.

Why did RORγt^-/-^ Th1 cells not induce EAE despite entering the CNS? It is tempting to speculate from our study that these cells did not survive well and long enough to produce inflammatory damage in the CNS or lacked an important pathogenic function. In this regard, we recently showed that iNOS activity during an immune response affects the CNS pathology in the brain and spinal cord in EAE (*45*). The presence of iNOS-expressing GFAP^+^ astrocytes in wild-type Th17 and RORγt^-/-^ Th1 infiltration indicated the possible role of iNOS-induced apoptosis of infiltrating immune cells in RORγt^-/-^ Th1 recipients. However, adoptively transferred RORγt^-/-^ Th1 cells failed to induce EAE in iNOS^-/-^ recipients, suggesting that iNOS- induced apoptosis of RORγt^-/-^ Th1 cells in the CNS parenchyma is dispensable for the induction of EAE. We then sought to determine the lack of pathogenic function in RORγt^-/-^ Th1 cells. The RORγt regulates the expression of several inflammatory molecules, such as IL-17A, IL- 17F, IL-21, IL-21R, IL-22, IL-23R, and GM-CSF in Th17 cells (*10, 11*). Our results showed that Th17 cytokines such as IL-17A, IL-21, and GM-CSF promote IL-23R expression in neurons, and these cytokines and their receptors are known to contribute to Th17-mediated neuronal autoimmunity (*13, 18, 29, 32, 33*). We show that Th1 cytokines such as IFN-γ or IL- 1β did not significantly induce the IL-23R expression in neuronal cells. It has been reported that neutralization or deficiency of IFN-γ does not prevent the EAE but exacerbates the disease (*45, 46*). Our observation of the presence of IL-23R-expressing NeuN^+^ and β-III tubulin^+^ neurons in the brain and spinal cord, respectively, of the MOG-immunized mice, drove us to investigate the role of IL-23-IL-23R signaling in neurons during EAE. Previously, neurons were shown to express IL-23R under oxygen and glucose deprivation conditions (*47*). Our data showed that the recipients of MOG-specific wild-type Th17 cells also showed IL-23R- expressing neurons in the CNS. However, the recipients of the RORγt^-/-^ Th1 cells did not show IL-23R expression on these neurons. The absence of IL-23R expression on other cell types of the inflamed CNS, such as astrocytes, microglial cells, and oligodendrocytes, suggests that cells other than Th17 cells and CD11b^+^ myeloid cells, neurons are also an important target of IL-23 signaling, which has not been explored clearly in the context of CNS autoimmunity.

Our data further show that IL-23 stimulation of IL-23R-expressing neurons undergoes apoptosis, as seen by increased cleaved caspase 3 localization in IL-23R^+^β-III tubulin^+^ neurons in the recipients of wild-type Th17 cells. A similar observation is missing in the RORγt^-/-^ Th1 cell recipients. The induction of IL-23R on neurons upon injections of purified recombinant murine IL-23 in the ventricles of the mice showed that IL-23 signaling has an autoregulatory function. The same phenomenon was not observed in the RORγt^-/-^ hosts as in RORγt^+/-^ mice and *i*.*c*.*v*. IL-23 injection-induced apoptosis in them. These observations lead us to think that IL-23R signaling in neurons appeared pathogenic and contributed to neuronal damage and the development of clinical EAE. The IL-23 is also shown to induce the apoptosis of cardiomyocytes and promote myocardial ischemia/reperfusion injury in rats (*48*). Furthermore, over-expression of IL-23R in 293ET and HeLa cells has been shown to activate caspase 3 and 9, which are critical for inducing apoptosis (*49*). Our data with murine ESCs-derived neural cells and Neuro-2a cells show that neurons express functional IL-23R on their surface, which upon IL-23 stimulation, induce the phosphorylation of STAT3 at Ser727 and Tyr705. The stimulation of IL-23/IL-23R signaling in these cells promoted increased levels of cleaved caspase 3 and cleaved PARP-1. This phenotype was completely ablated in the cells that lacked IL-23R expression, suggesting that the IL-23/IL-23R signal drove the apoptotic program in the neurons. Cleaved caspase 3 is a key executor of cell-intrinsic, and extrinsic apoptosis pathways and an important inducer of PARP-1 cleavage, both of these events boost the apoptosis cell death process (*36, 37*). Furthermore, the increased frequency of Annexin-V and 7-AAD double-positive cells among these IL-23 treated cells shows increased late-phase apoptosis event, suggesting that IL-23 signaling via STAT3 activation might contribute to the apoptosis of neurons involving the activation of caspase 3 and cleavage of PARP-1. While it became clear that IL-23-IL-23R signaling induced the apoptotic death of the neurons, what triggers IL- 23R expression on the neuron during EAE is unknown. Our data with *in vitro* neuronal cell culture revealed that purified GM-CSF, TNF-α, IL-6, IL-17A, and IL-21 could induce IL-23R on the surface of neurons. Various cell types known to produce these pro-inflammatory cytokines are present in the active EAE lesions and multiple areas of the inflamed CNS, including highly pathogenic Th17 cells that produce IL-17A, GM-CSF, and IL-21 other cytokines (*10, 11*). Such an inflammatory milieu likely sets the neurons responsive to IL-23- IL-23R signaling leading to damage to the subsets of neurons (at least MAP2^+^, bIII-tubulin^+^) during EAE.

These findings highlight the differential requirement of T-bet and RORγt in Th1 and Th17 cells to promote neuroinflammation at BBB and CNS parenchyma levels in EAE. The present study gave a mechanism wherein neurons were forced to induce IL-23R expression when potently encephalitogenic T cells entered the CNS and set the inflammation in the CNS, and IL-23 locally produced by infiltrated activated DCs/macrophages glial cells, triggering apoptotic program in the IL-23R-expressing neurons. Understanding these events in greater detail would pave the way to finding therapeutic strategies to control neuronal inflammation and autoimmunity.

## Material and methods

### Mice

C57BL/6, congenic CD45.1 (PTPRC), T-bet^-/-^ (B6.129S6-Tbx21^tm1Glm^/J) and RORγt^-/-^ (B6.129P2(Cg)-Rorc^tm2Litt^/J) mice were purchased from The Jackson Laboratory (Bar Harbor, ME). The mice were bred and maintained in the Experimental Animal Facility of the National Centre for Cell Science (NCCS), Pune. All animal experiments were performed as per approved protocols from the Institutional Animal Ethics Committee (Project number: EAF/2010/B-258 and EAF/2016/B-259).

### Induction of passive EAE

Wild-type C57BL/6, T-bet^-/-^ or RORγt^-/-^ mice were subcutaneously (s.c.) injected with myelin oligodendrocytes peptide (MOG_35-55_; 200 μg/mouse) emulsified in complete Freund’s adjuvant (CFA) containing *Mycobacterium tuberculosis H37Ra* (5 mg/ml). After ten days, spleen and draining lymph nodes were harvested, and single-cell suspensions were prepared. Cells were differentiated into Th1 or Th17 in the presence of MOG peptide (50 μg/ml) in the RPMI-1640 medium supplemented with 10% FBS, 1 mM sodium pyruvate, 2 mM L-glutamine, 1X non-essential amino acids, 2 X 10^-5^ M 2-mercaptoethanol, 100U/ml penicillin and 100 μg/ml streptomycin along with either recombinant mouse IL-12 (10 ng/ml), anti-IL-4 (10 μg/ml) (for Th1 lineage) or IL-23 (10 ng/ml), IL-1ϕ3 (10 ng/ml), anti-IFN-γ (10 μg/ml) and anti-IL-4 (10 μg/ml) (for Th17 lineage) at 37^°^C for four days. Live cells were isolated by Histopaque 1083 (Sigma) density-gradient centrifugation, and 20 X 10^6^ cells/mouse was adoptively transferred into sub-lethally irradiated (350 rads) congenic CD45.1 mice. Pertussis toxin (200 ng/mouse) was given intravenously (i.v.) on days 0 and 2. The mice were observed daily for the development of clinical symptoms, and mice were clinically scored as- score 0, no symptoms; 1, limp tail or hind limb weakness, but not both; 2, both limp tail and hindlimb weakness; 3, complete hind limb paralysis; 4, forelimb paralysis; and 5, death.

### Induction of active EAE

Active EAE was induced by subcutaneous (*s.c*.) injection of MOG_35-55_ (MOG) peptide (200 μg/mouse) emulsified in complete Freund’s adjuvant (CFA) containing *Mycobacterium tuberculosis H37Ra* (5 mg/ml) at the flanks of mice. Pertussis toxin (200 ng/mouse) was given intravenously (*i.v.*) on days 0 and 2. The mice were observed daily for the development of clinical symptoms, and mice were clinically scored.

### Adoptive transfer of CD4^+^ T cells

Naive CD4^+^CD25^-^ T cells from wild-type C57BL/6, T-bet^-/-^ or RORγt^-/-^ mice were purified using flow cytometry sorting. Purified CD4^+^CD25^-^ T cells (2 x 10^6^ cells/mouse) were adoptively transferred into T-bet^-/-^ mice. Subcutaneous injection of MOG (200 μg/mouse) peptide emulsified in complete Freund’s adjuvant (CFA) containing *Mycobacterium tuberculosis H37Ra* (5 mg/ml) was injected at the flanks of mice, and also *i.v.* Pertussis toxin (200 ng/mouse) was given on days 0 and 2. The mice were monitored for the development of clinical symptoms of EAE and clinically scored as described above.

### Immunofluorescence staining of the brain and spinal cord

Mice were sacrificed and transcardially perfused with ice-cold PBS, and the brain and spinal cord were harvested and immediately snap-frozen in the OCT compound (Sakura Finetek, Torrance, CA). Eight-micrometer-thick cryosections were fixed in chilled acetone for 5 minutes, air dried, washed with PBS, and blocked with 10% horse serum (Jackson ImmunoResearch, West Grove, PA) for 30 minutes at room temperature (RT). Sections were washed three times with ice-cold PBS and incubated with primary antibodies (1:200 dilutions) overnight (12-14 hours) at 4°C in the dark, followed by washed four times with ice-cold PBS and incubated with fluorochrome-conjugated secondary reagents (1:1000 dilution) for 60 minutes at RT in the dark. Sections were washed four times with PBS, fixed with 1% paraformaldehyde (PFA) for 5 minutes at RT, and mounted with DAPI-containing mounting medium (Electron Microscopy Sciences, Hatfield, PA). Sections were visualized under Leica DMI6000 inverted fluorescent microscope (Leica Microsystems, Germany) or Zeiss LSM510 META confocal microscope (Carl Zeiss, Germany). Images were analyzed using Leica MMAF software (Leica) or Image-J software (NIH, USA).

### Intracellular cytokine staining

Mice were sacrificed at the indicated time points, spleen and lymph nodes were harvested, made single-cell suspensions, and removed RBCs using ACK lysis buffer. Cells were washed with RPMI-1640 used for intracellular cytokine staining. Cells (6 X 10^6^ cells/well) were stimulated in 24-well plates with phorbol-myristate acetate (PMA; 50 ng/ml) and ionomycin (850 ng/ml) in the presence of brefeldin-A and monensin (2 μM) at 37^°^C in a humidified 5% CO_2_ incubator for 6 hours. Cells were harvested, washed with PBS, and stained with APC-anti-mouse CD4 on ice for 30 minutes. Cells were washed with PBS, fixed, and permeabilized using a Foxp3 fixation/permeabilization buffer kit (BioLegend). The cells were stained with Brilliant Violet 421 anti-mouse IL-17A and PE/Cy7 anti-mouse IFN-γ antibodies. Finally, cells were washed, fixed in 1% PFA, and acquired using flow cytometry (FACS Canto II, BD Biosciences) and analyzed using Flowjo software (Tree Star).

### Intrathecal (i.t.) injection of IFN-γ

Stereotactic intrathecal (*i.t.*) injections in mouse cortices were performed as described earlier (Argaw et al., 2012). T-bet^-/-^ mice (12 weeks old) immunized with 200 μg MOG in CFA emulsion were given i.t. IFN-γ injection at day 15. Briefly, T-bet^-/-^ were anesthetized using ketamine/xylazine and given stereotactic *i.t.* injection of purified recombinant IFN-γ (50 pg/mouse) in two microliters of PBS at a rate of 0.25 μl/minute. Age- and sex-matched control mice were given a vehicle (2.0 μl PBS) at the same rate. The coordinates used for *i.t.* injection was; 1 mm posterior, 2 mm left lateral, and 1.5 mm dorsoventral to the Bregma. Mice skulls were applied with antiseptic lotion, and the skin over the skull was stitched. Food and water were provided on the flooring of animal cages. After three days, the mice were transcardially perfused with ice-cold PBS. The brain and spinal cord were harvested and snap-frozen in OCT. Eight micrometer-thick cryo-sections were made and stained for multicolor immunofluorescence microscopy.

### Quantitative real-time PCR

RNA was isolated from cultured Neuro-2a cells using TRIzol reagent (Invitrogen, Carlsbad, CA), cDNAs prepared using oligo d(T)_12-16_ primers (Invitrogen). The mRNA expressions of the specific gene were quantified by the CFX96 thermal cycler (Biorad, Hercules, CA) using the SYBR Green PCR kit (Life Technologies, Carlsbad, CA). PCR condition consisted of at 50^0^C for 2 minutes, at 95^0^C for 10 minutes, at 95^0^C for 30 seconds, and at 60^0^C for 30 seconds. The relative mRNA expression of a gene was calculated as 2^(Ct^ ^of^ ^control gene – Ct of a specific gene)^. The sequences of primers used are - IL-23R (forward 5’- TGCCATGACTCCAGGATGGAACTT-3’); reverse (5’- GCAAACAAGCAAACAAGCACAGCC-3’), GAPDH (forward 5’- GACATGCCGCCTGGAAGAAAC-3’); reverse (5’-AGCCCAGGATGCCCTTTAGT-3’).

### Intracerebroventricular (i.c.v.) injection of IL-23

Stereotactic intracerebroventricular (*i.c.v.*) injections were performed as described earlier (DeVos and Miller, 2013). Briefly, RORγt^+/-^ or RORγt^-/-^ mice (10-12 weeks old) were anesthetized using ketamine/xylazine and given stereotactic *i.c.v.* injection of purified recombinant mouse IL-23 (200 pg/mouse) in ten microliters of PBS at a rate of 2.0 μl/minute. The coordinates used for *i.c.v.* injections were; 0.3 mm anterior, 1.0 mm right lateral, and 3.0 mm dorsoventral to the Bregma. Mice skulls were applied with antiseptic lotion, and the skin over the skull was stitched. Food and water were provided on the flooring of animal cages. After three days, the mice were transcardially perfused with ice-cold PBS. The brain and spinal cord were harvested and snap-frozen in OCT. Eight-micrometer-thick cryosections were made and stained for multicolor immunofluorescence microscopy.

### Differentiation of neuronal cells from embryonic stem cells

Murine embryonic stem (ESCs) line D3 was procured from ATCC and was maintained in the undifferentiated state. ESCs were differentiated into neural lineage following the monoculture strategy described earlier (*50*). In brief, ESCs were cultured using the maintenance medium [DMEM with 15 % FBS (Hyclone), L-glutamine, penicillin-streptomycin, non-essential amino acids (all from Invitrogen), β-mercaptoethanol (Sigma-Aldrich) and leukemia inhibitory factor (LIF, Chemicon)], and passaged every 48 hours. For neural differentiation in monoculture, the cells were dispersed and seeded on gelatin-coated dishes using DMEM with 10% FBS and other additives except for LIF. Subsequently, the cells were maintained under serum replacement conditions (*50*). Considering the time window for neural progenitor generation to be 6-9 days, as reported earlier, and day 14 showed optimum neuronal differentiation (∼80% Map2^+^ cells) (*50*), we used day 14 cultured cells to assess IL- 23 receptor expression and apoptosis of neural cells.

### Knockdown of IL-23R in Neuro-2a cells

Neuro-2a cells were infected with a lentivirus vector encoding 19-25nt (plus hairpin) IL-23R shRNA (Santa Cruz Biotechnology; sc-60835-SH) with 8 μg/ml polybrene. After 72 hours of infection, IL-23R knockdown Neuro-2a cells were selected using increasing concentration (1-5 μg/ml) of Puromycin. After three passages in the puromycin selection media, expression was confirmed using Western blot and used for the experiments.

### Apoptosis of Neuro-2a cells

Neuro-2a cells (2 X 10^6^ cells/well) were cultured in 6-well plates in complete Eagle’s Minimum Essential Medium. Cells were untreated or treated with recombinant mouse IL-23 (10 ng/ml) at 37^0^C in a humidified 5% CO_2_ incubator for 48 hours. The cells were harvested and surface stained for IL-23R, fixed in 4% paraformaldehyde (PFA), and permeabilized with 0.05% saponin-containing PBS. The cells were stained for intracellular cleaved caspase 3. Further, apoptosis of the Neuro-2a cell was analyzed by Annexin-V-FITC and 7-AAD staining accordingly to the manufacturer’s instruction (BioLegend). Cells were immediately acquired on a Canto II flow cytometer (BD Biosciences). For immunofluorescence analysis, Neuro-2a cells (2.5 X 10^4^ /well) grown on glass-coverslip in 12 well plates, fixed with 4% PFA, permeabilized with 0.5% Triton-X 100, blocked with 5% BSA and 5% normal horse serum in PBST (PBS + 0.05% Tween 20), and incubated with primary antibodies overnight at 4°C, followed by five washes with PBST. Cells were stained with secondary antibodies at RT for 60 minutes in the dark, followed by six times washed with PBST and nuclear stained with DAPI. Cells were fixed with 1% PFA and mounted in a mounting medium containing Tris buffer. Images were acquired using a Leica DMI6000 inverted fluorescent microscope (Leica Microsystems, Germany), and data were analyzed using Leica MMAF software (Leica).

### Western Blotting

Neuro-2a cells were treated with IFN-γ, GM-CSF, and IL-1β at 20 ng/ml for indicated time points, washed with ice-cold PBS, and lysed in RIPA buffer containing 50 mM Tris.HCl pH 8.0, 150 mM NaCl, 1% NP-40, 0.1% SDS, 1 mM DTT, protease inhibitor cocktail, and 1 mM PMSF. Cell debris was removed by centrifugation, and protein concentration in the supernatant was estimated using the Bradford reagent. About 30 μg protein/well was resolved using SDS-PAGE. Proteins were transferred onto the PVDF membrane. The membrane was blocked with 5% non-fat milk at room temperature for 60 minutes. The membrane was incubated with primary antibodies at 1:2000 dilution at 4℃ overnight. Membranes were washed and incubated with HRP-conjugated secondary antibodies (1:6000) at room temperature for 60 minutes. Rabbit anti-Cyclophilin B (1:2000; Abcam, Cambridge, MA) was used as a loading control. The blot was developed using a chemiluminescent ECL system as per the manufacturer’s guidelines (ThermoFisher Scientific). The band intensity was analyzed using NIH ImageJ software.

## Limitations of the study

In this study, we did not use neurons from IL-23R^-/-^ mice to characterize the apoptosis and the source of immune cells for IL-23 in the CNS induced by RORγt^+^ CD4 T cells. We used IL-23R siRNA to downregulate the IL-23 receptors in neurons and treated them with IL-23 in vitro to substantiate ours *in vivo* findings. Although, it is well known that IL-23p19^-/-^ and IL- 23R^-/-^ mice do not show demyelination in the CNS with MOG peptide injection and are protected from EAE (*17, 18*).

### Statistical analyses

Statistical analysis was performed using GraphPad Prism 6 software (GraphPad, San Diego, CA). The unpaired two-tailed Student *t*-test was used, assuming that data were normally distributed and had an equal variance between the groups under comparison. Other statistical methods used are described in the figure legends. A *p*<0.05 was considered statistically significant.

## Acknowledgments

We thank Ramanamurthy Boppana and Rahul Bankar for their help with animal experiments. SAS received Senior Research Fellowship from the Council of Scientific and Industrial Research, and THM received INSPIRE fellowship from the Department of Science and Technology, Government of India. THM received GL received NCCS intramural funding and grants from the Department of Biotechnology (Grants numbers, BT/PR4610/MED/30/720/2012, BT/RLF/Re-entry/41/2010, BT/PR14156/BRB/10/1515/2016), Department of Science and Technology (DST/SJF/LSA- 01/2017-18), Government of India.

## List of Supplementary Materials

**A. Supplementary Figures**

Figure S1. T-bet^-/-^ Th17 cells do not alter the localization of claudin-5 on BBB during EAE.
Figure S2. Effect of cytokine stimulation of neuronal cells on IL-23R expression.
Figure S3. Embryonic stem cells-derived neuronal cells express IL-23R and undergo apoptosis after stimulation with IL-23.

**B. Reagents detail.**

## Abbreviations

BBB: blood-brain barrier
CNS: central nervous system
DAPI: 4’,6- diamidino-2-phenylindole
EAE: experimental autoimmune encephalomyelitis
MOG: myelin oligodendrocyte glycoprotein peptide_35-55_
MS: multiple sclerosis
PTx: pertussis toxin
RORγt: RAR-related orphan receptor gamma t
T-bet: T-box transcription factor
TEM: transendothelial migration
VE-cadherin: vascular endothelial-cadherin
ZO-1: zona occludens-1.

## Supplementary figure legends

**Figure S1.**
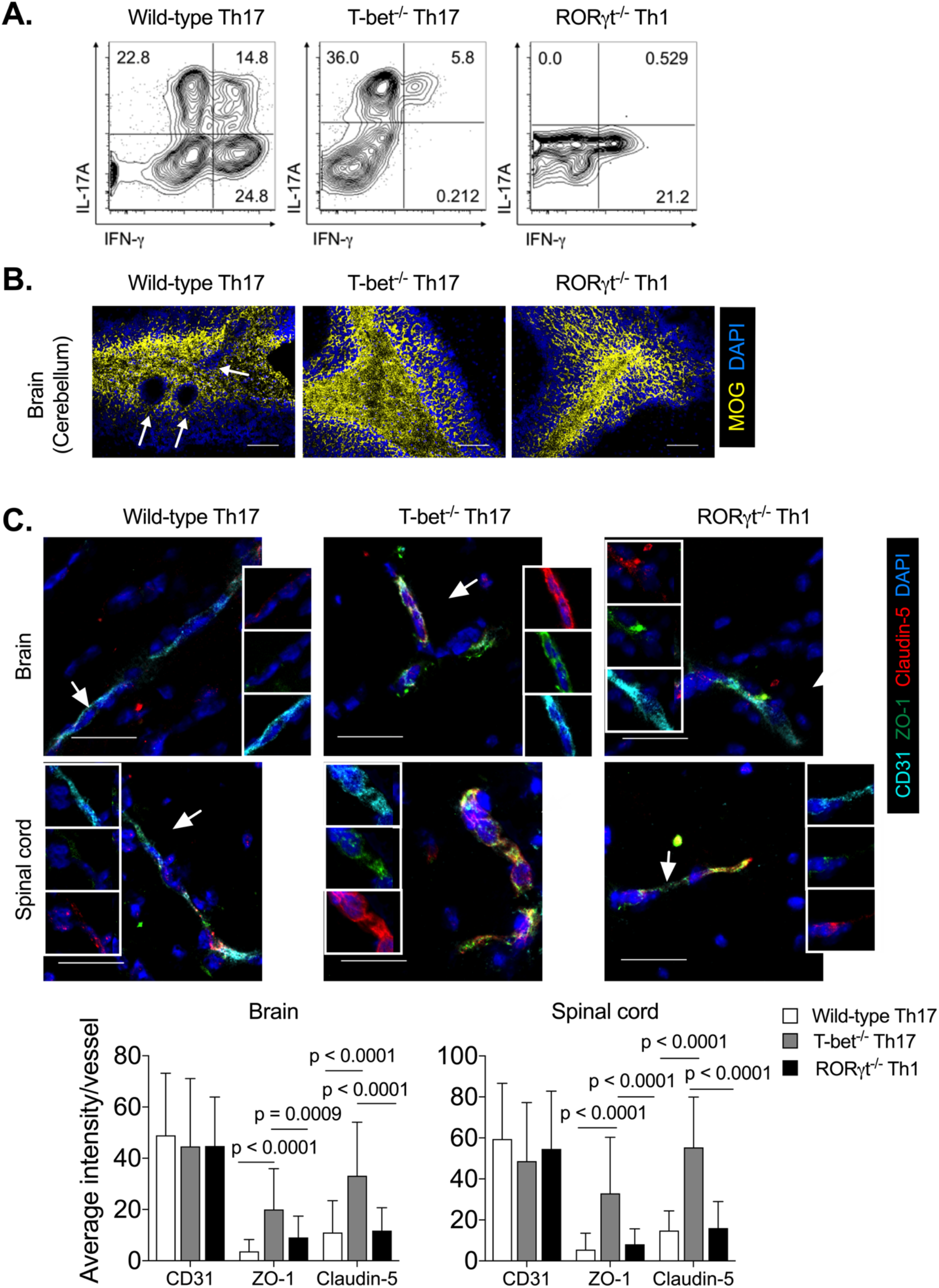
T-bet^-/-^ Th17 cells do not alter the localization of claudin-5 on BBB during EAE. **(A)** In-vitro differentiated MOG-specific CD4^+^ T cells were stimulated with PMA and Ionomycin in the presence of Brefeldin-A for 5 hours, and intracellular cytokine expression was analyzed. Dot plots show IL-17A and IFN-γ expressing CD4^+^ T cells gated on activated CD4^+^CD25^+^ T cells. Numbers in the gates represent percentages of cytokine-producing cells. **(B)** The brain cryo-section of mice, as in Figure 1A, was stained with MOG (yellow) and nuclear stain DAPI (blue), and the representative images are shown. Arrows in the images indicate the regions of focal lesion and demyelination. **(C)** The brain and spinal cord of mice from Figure 1A are stained with CD31^+^ (light blue), ZO-1 (green), claudin-5 (red), and nuclear stain DAPI (dark blue). The representative images are shown (upper). Quantitation of the average intensity of CD31, ZO-1, and claudin-5 per vessel is shown (lower). A total of 46 to 75 vessels in the brain and about 49 to 55 vessels in the spinal cord were analyzed. Data are representative of two independent experiments with 3-5 mice brain and spinal cord tissues from each group. Statistical significance between the groups is determined by One-way ANOVA followed by Tukey’s test **(C)**; **the** Error bar represents S.D. Original magnification 200x **(B),** 630x **(C)**; Scale bar 100μm **(B)** and 30μm **(C)**.

**Figure S2.**
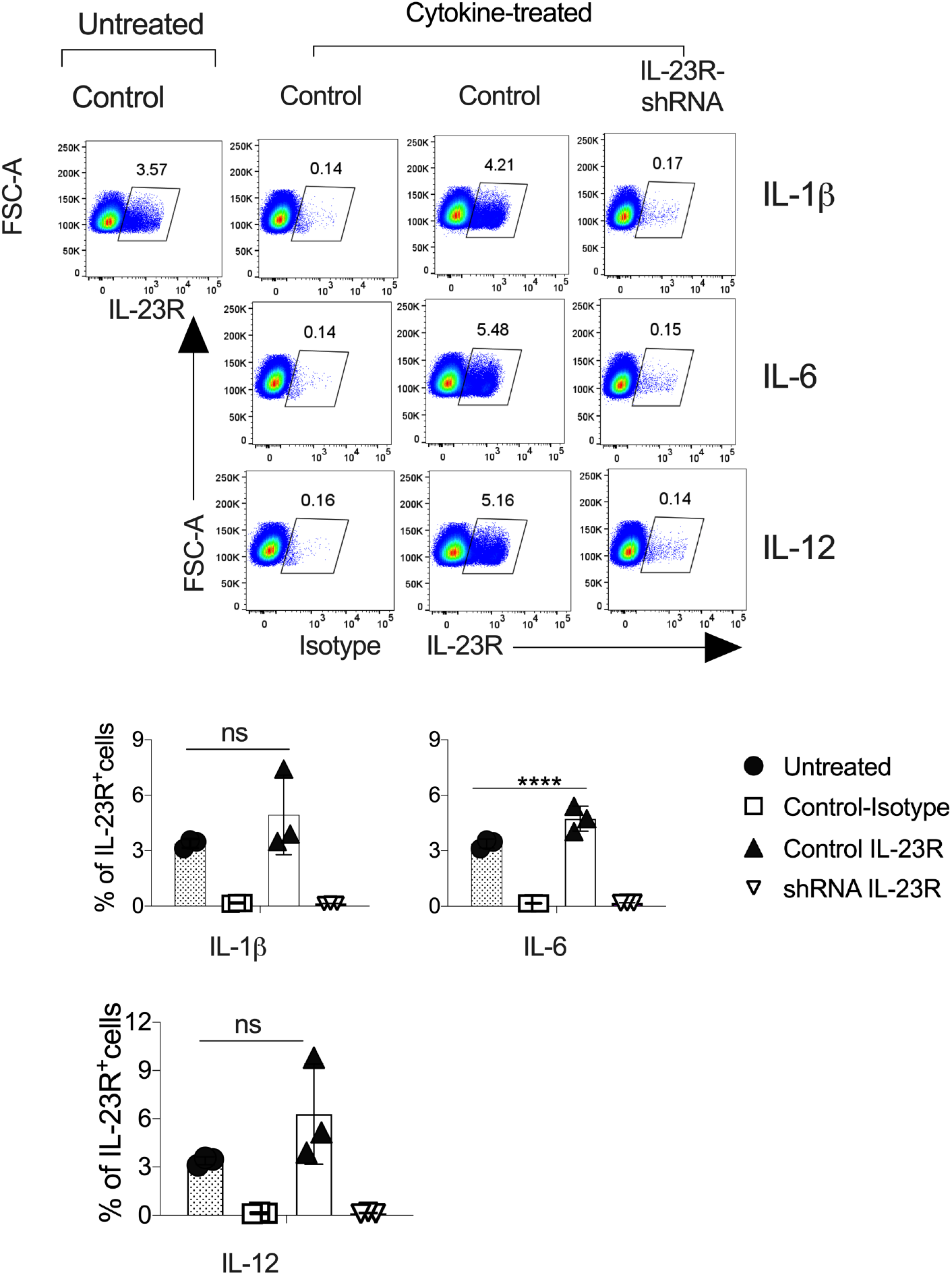
Effect of cytokine stimulation of neuronal cells on IL-23R expression. Neuro- 2a cells (control cells) or IL-23R-knockdown Neuro 2a cells (IL-23R-shRNA) were treated with recombinant purified IL-1β, IL-6, or IL-12 (10 ng/ml) for 48 hours, and IL-23R expression was monitored using flow cytometry. A representative dot plot is shown (top). Each symbol represents data from an individual test. The error bar represents SEM. Statistical significance between control neuro2a cells treated and untreated samples are determined by One-way ANOVA. ns = not significant. ****p>0.0001.

**Figure S3.**
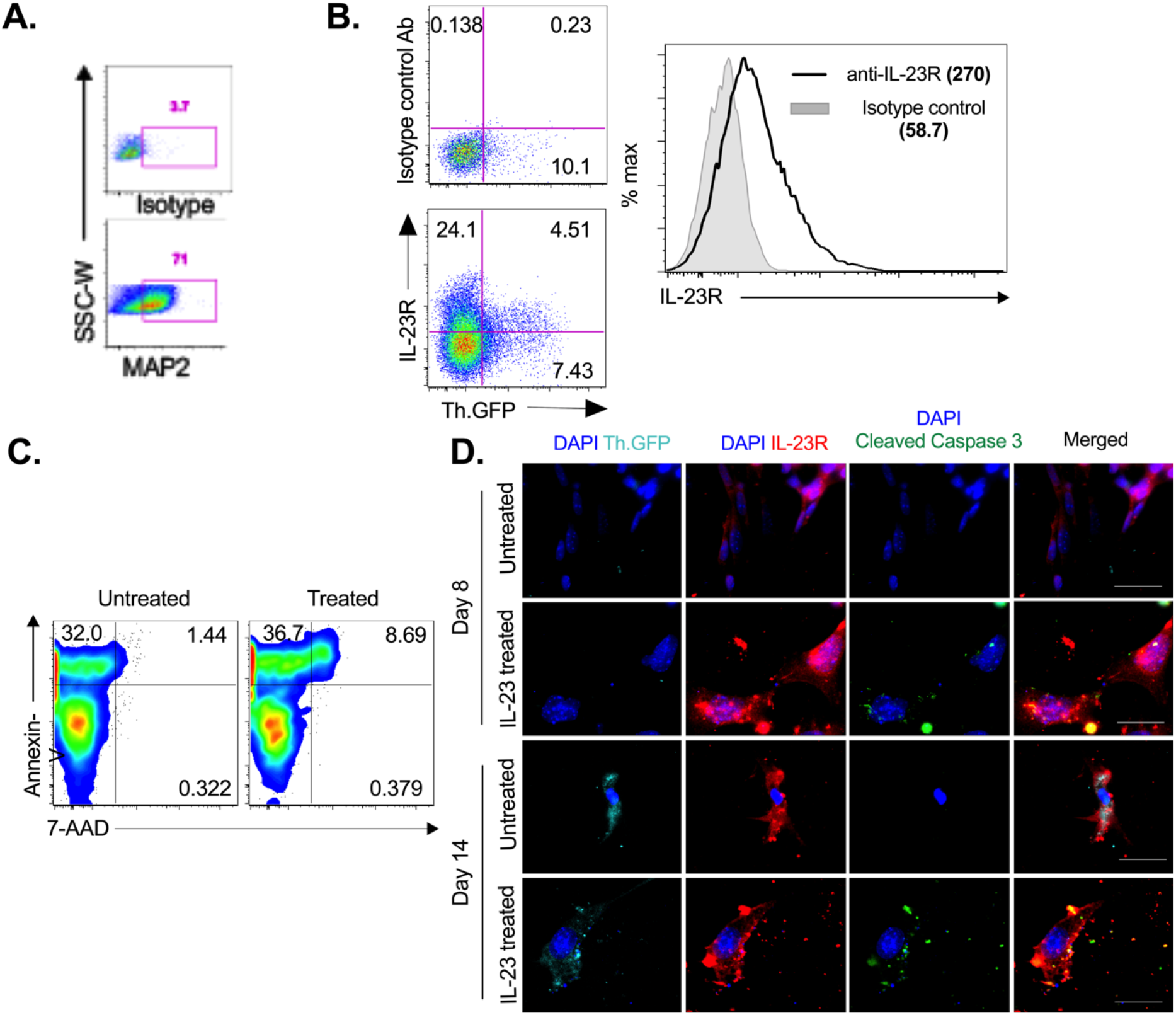
Embryonic stem cells-derived neuronal cells express IL-23R and undergo apoptosis after stimulation with IL-23. Embryonic stem cells (ESCs) were differentiated into the neural lineage. On day 14, Neuronal cells were characterized and used for experiments. **(A)** The surface expression of neuronal marker MAP2 was analyzed and plotted. **(B)** Dot plots show the surface expression of IL-23R on differentiated Th.GFP-expressing neurons and histograms show the mean fluorescence intensity (MFI) of surface IL-23R on total neuronal cells. **(C)** Neuronal cells were stimulated with purified IL-23 for 48 hours, and apoptosis of neurons was measured with Annexin-V and 7-AAD. Numbers in the gates show the percentage of indicated cell populations**. (D)**. *N*euronal cells were treated with IL-23 (10 ng/ml) for the last 48 hours. Cells were fixed and stained for IL-23R, cleaved caspase 3, and nuclear stain DAPI. Representative images of IL-23R (red), tyrosine hydroxylase-GFP (light blue), cleaved caspase 3 (Green), and DAPI (dark blue) were shown. Original magnification 1000x, scale bar 30 μm. The data shown represent two **(C, D)** to three **(A, B)** independent experiments.

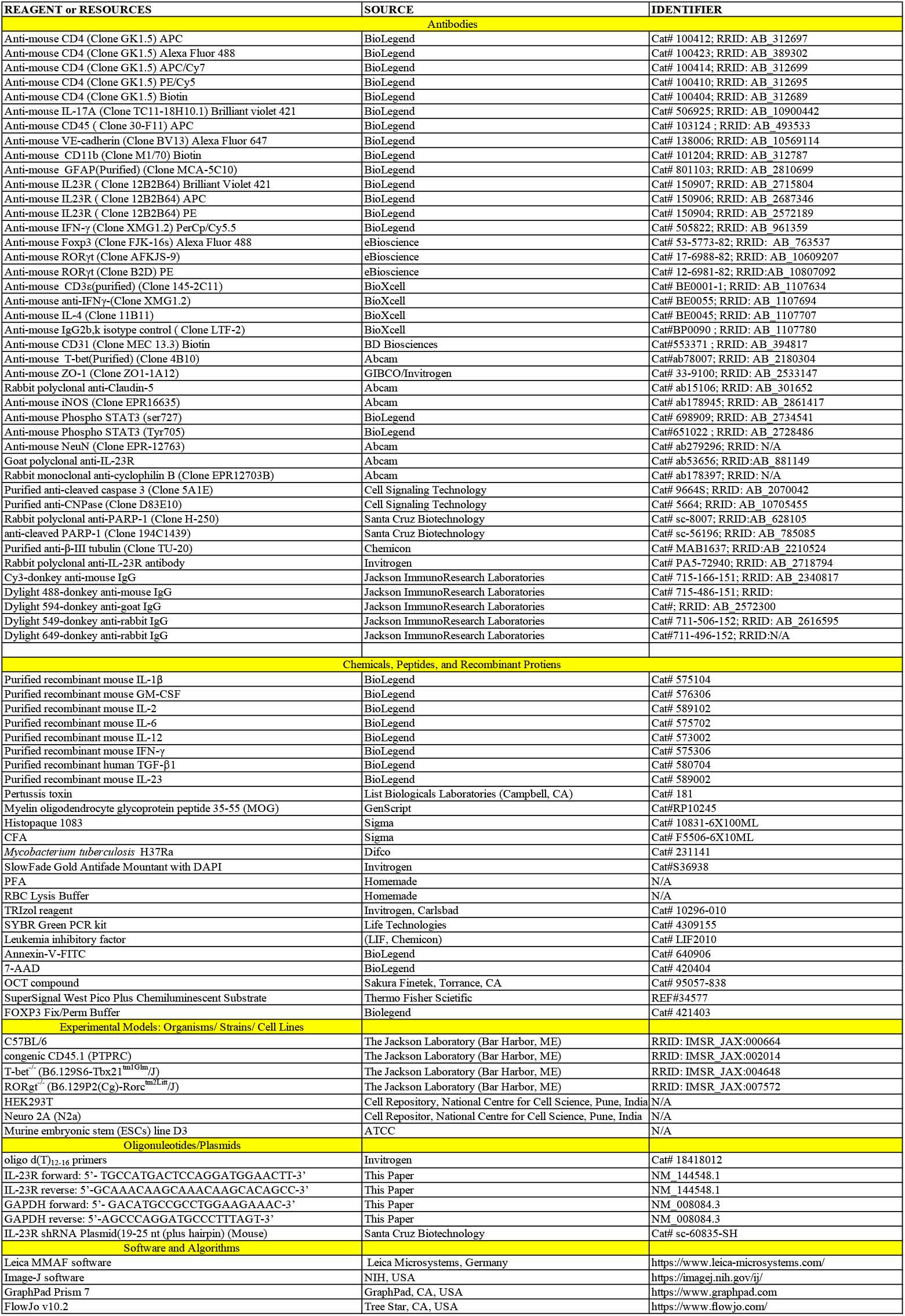

